## Supplemental materials for "High-throughput analysis of single human cells reveals the complex nature of DNA replication timing control"

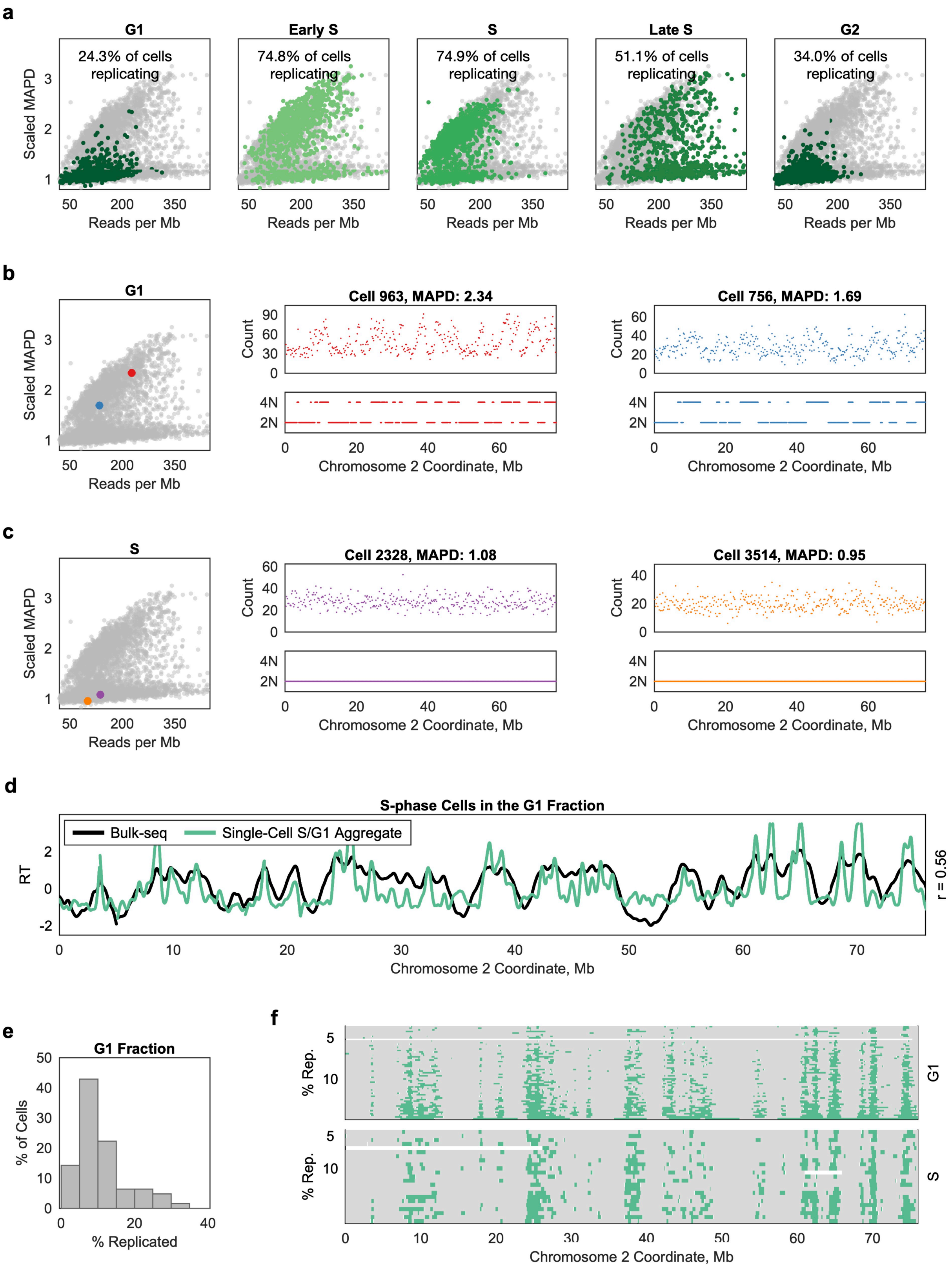


Supplementary Figure 1. ***In silico* sorting of cells recapitulates fluorescence activated cell sorting (FACS).**

**a** GM12878 cells were sorted into five fractions by FACS prior to sequencing. For each fraction, the corresponding cells are highlighted in green. Each fraction contains a mixture of $G_1$/$G_2$- and S-phase cells according to *in silico* sorting (as in Figure 1b). **b** The FACS $G_1$ fraction contained several cells with higher scaled MAPD than expected for a non-replicating cell. Two copy-number states were inferred across each chromosome, suggesting that these cells were replicating. **c** The FACS S-phase fraction contained many cells with lower scaled MAPD than expected for a replicating cell. A sole copy-number state was inferred across each chromosome, suggesting that these cells were not replicating. **d** An S/$G_1$ aggregate replication timing profile inferred from the $G_1$ FACS fraction library, based on preliminary *in silico* assignment of 323 cells as "S", recapitulated the bulk-sequencing profile for this cell line. **e** Cells within the $G_1$ FACS fraction assigned as replicating after copy-number inference (n = 63) were less than 35% replicated (*i.e.*, in early S phase). There was one outlier (87% replicated). **f** Replication profiles for early S-phase cells in the $G_1$ and S FACS fractions were visually indistinguishable.

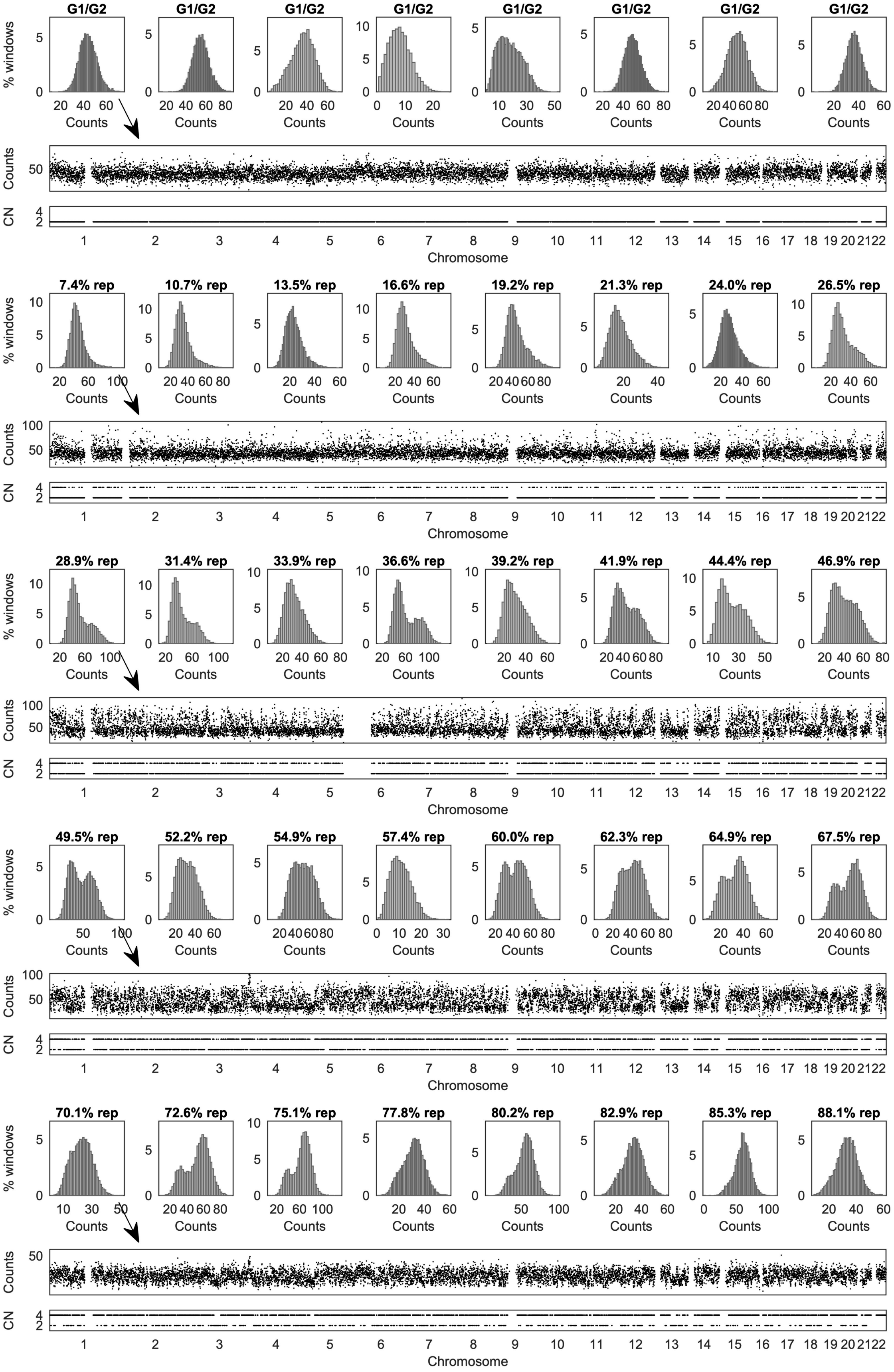

G1/G2
7.4% rep
10.7% rep
13.5% rep
16.6% rep
19.2% rep
21.3% rep
24.0% rep
26.5% rep
28.9% rep
31.4% rep
33.9% rep
36.6% rep
39.2% rep
41.9% rep
44.4% rep
46.9% rep
49.5% rep
52.2% rep
54.9% rep
57.4% rep
60.0% rep
62.3% rep
64.9% rep
67.5% rep
70.1% rep
72.6% rep
75.1% rep
77.8% rep
80.2% rep
82.9% rep
85.3% rep
88.1% rep
% windows
Counts
CN
Chromosome
1 2 3 4 5 6 7 8 9 10 11 12 13 14 15 16 17 18 19 20 21 22

Supplementary Figure 2. **Distribution of single-cell read counts in 200kb windows depends on S-phase progression.** Forty single cells from the S-phase FACS fraction library are shown. Read counts are distributed around a single mode in cells in $G_1$/$G_2$ (top row). The bimodal distribution of read counts in replicating cells is most apparent in mid-S phase. For the left-most cell in each row, read counts and corresponding copy-number inferences (CN) are displayed for the whole genome. Representative examples were selected to reflect patterns across S-phase progression.

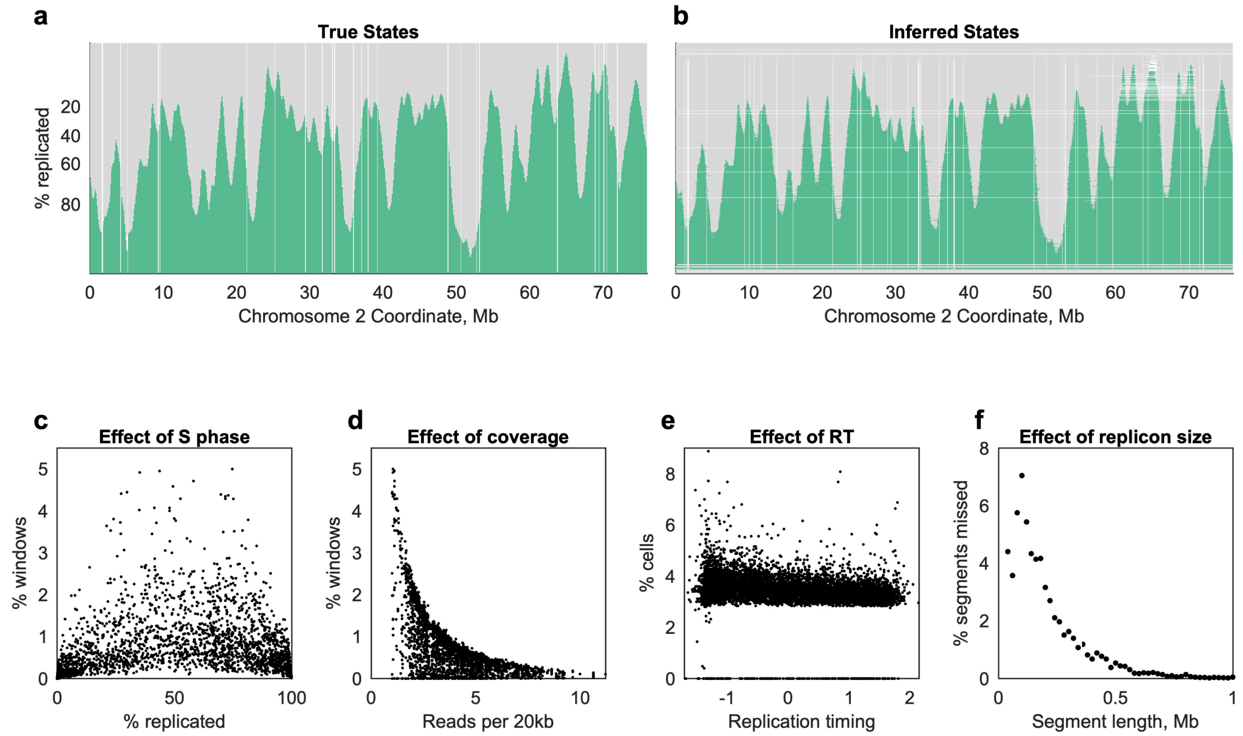


Supplementary Figure 3. **Assessment of replication state inference by hidden Markov model (HMM) with simulated data.** **a** 2,500 cells were simulated with known "true" replication states at each genomic window, distributed across S phase. Each row represents one simulated cell. **b** Raw read counts were drawn from a Poisson process for each "true" replication state, with rate coefficients chosen to match the coverage of the GM12878 unsorted library. These raw counts were then processed with the same pipeline as the observed data. Each row represents the corresponding cell in **a**. Incorrect copy-number inferences were most frequent in mid-S phase cells (**c**) and in lower-coverage cells (**d**), while the effects of replication timing were mild (**e**). **f** Copy-number inference allowed identification of 98.6% of "true" replicons. Replicons comprised of a single 20kb window were frequently missed (26.8% not successfully detected). Replicons comprised of at least two windows were detected in >90% of cases.

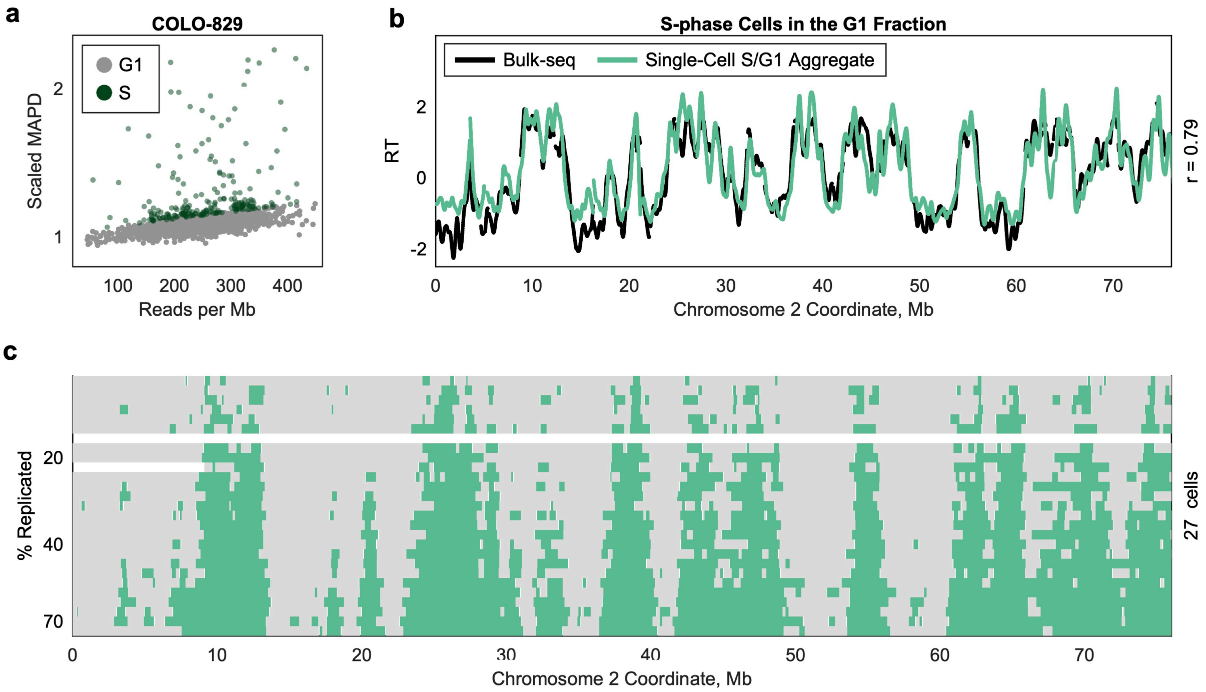


Supplementary Figure 4. **S-phase contamination was observed in published $G_1$ FACS data**. **a** *In silico* sorting of 1,475 cells from the human melanoma cell line COLO-829 identified 233 potentially replicating cells in an experiment where FACS was used to isolated only $G_1$ cells for copy-number aberration analysis[1]. **b** The aggregate S/$G_1$ replication timing profile inferred using *in silico* sorting assignments recapitulated the ensemble replication timing profile measured in COLO-829. **c** Replication profiles for 32 cells were consistent with being in S phase and demonstrated a uniform replication progression pattern. Five cells are not shown due to chromosome-specific copy number anomalies.

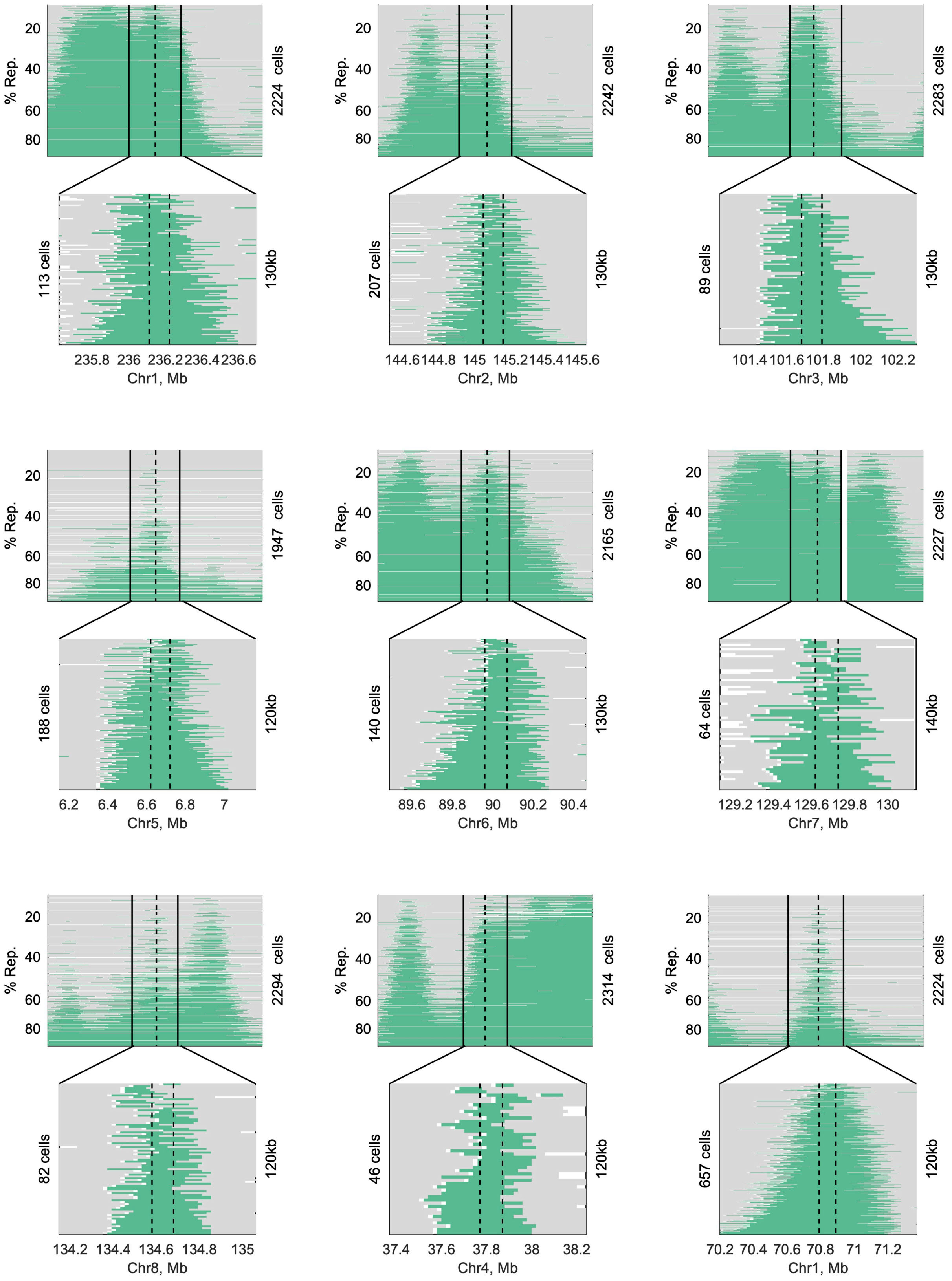
% Rep.
20
40
60
80
2224 cells
113 cells
130kb
235.8 236 236.2 236.4 236.6
Chr1, Mb
2242 cells
207 cells
130kb
144.6 144.8 145 145.2 145.4 145.6
Chr2, Mb
2283 cells
89 cells
130kb
101.4 101.6 101.8 102 102.2
Chr3, Mb
1947 cells
188 cells
120kb
6.2 6.4 6.6 6.8 7
Chr5, Mb
2165 cells
140 cells
130kb
89.6 89.8 90 90.2 90.4
Chr6, Mb
2227 cells
64 cells
140kb
129.2 129.4 129.6 129.8 130
Chr7, Mb
2294 cells
82 cells
120kb
134.2 134.4 134.6 134.8 135
Chr8, Mb
2314 cells
46 cells
120kb
37.4 37.6 37.8 38 38.2
Chr4, Mb
2224 cells
657 cells
120kb
70.2 70.4 70.6 70.8 71 71.2
Chr1, Mb

Supplementary Figure 5. **Broad initiation regions (*i.e.*, those wider than 120kb) often appear visually to contain multiple non-overlapping replication tracks.** Eight representative examples are shown, in which a single IR was called, but visual inspection suggests multiple discrete initiation sites that have not been disambiguated, supporting the notion that these IR widths are overestimated. One counterexample is shown in the bottom right (Chr1, ~70.8Mb) – this IR's width may be due to it encompassing a cluster of origins in close proximity or to replication fork asymmetry.

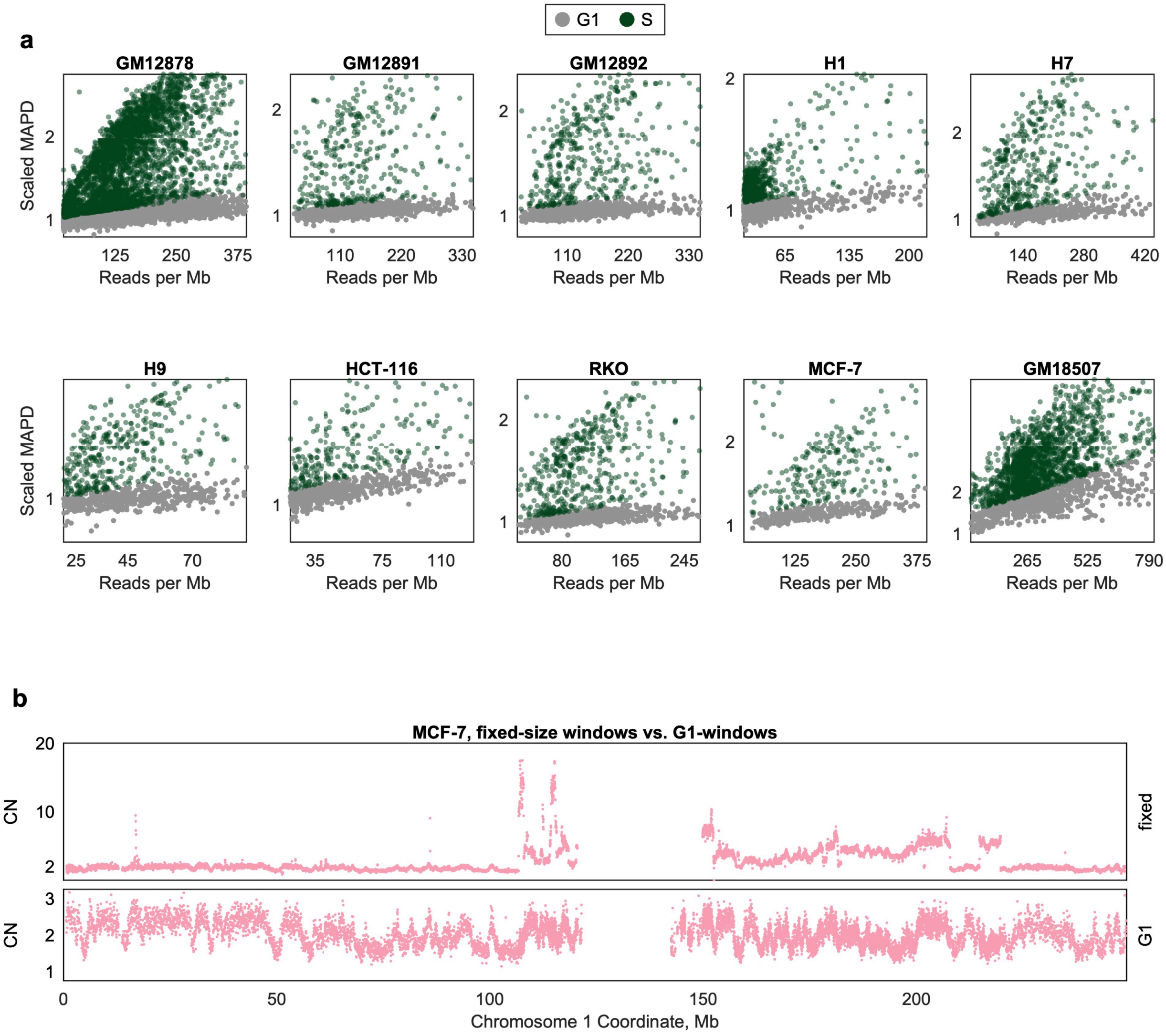


Supplementary Figure 6. **Metrics of *in silico* cell sorting for all cell lines under study. a** *In silico* sorting identifies two distinct populations of cells based on the relationship between mean reads per Mb and scaled MAPD, corresponding to $G_1$/$G_2$- and S-phase cells. GM18507 cells were sequenced after DLP+ library preparation[2]. **b** *In silico* sorting is robust in aneuploid cancer cell lines (*e.g.*, MCF-7) and corrects for the effects of copy-number aberrations (CNAs). Top panel: MCF-7 contains numerous CNAs with a wide range of copy-number values. Each dot represents the sum of read counts across all *in silico* assigned S-phase cells in fixed 20kb windows. Bottom panel: Using *in silico* assigned $G_1$/$G_2$ cells to define genome windows reveals more subtle copy-number fluctuations corresponding to replication timing. Each dot represents the sum of read counts across all *in silico* assigned S-phase cells in $G_1$-defined windows.

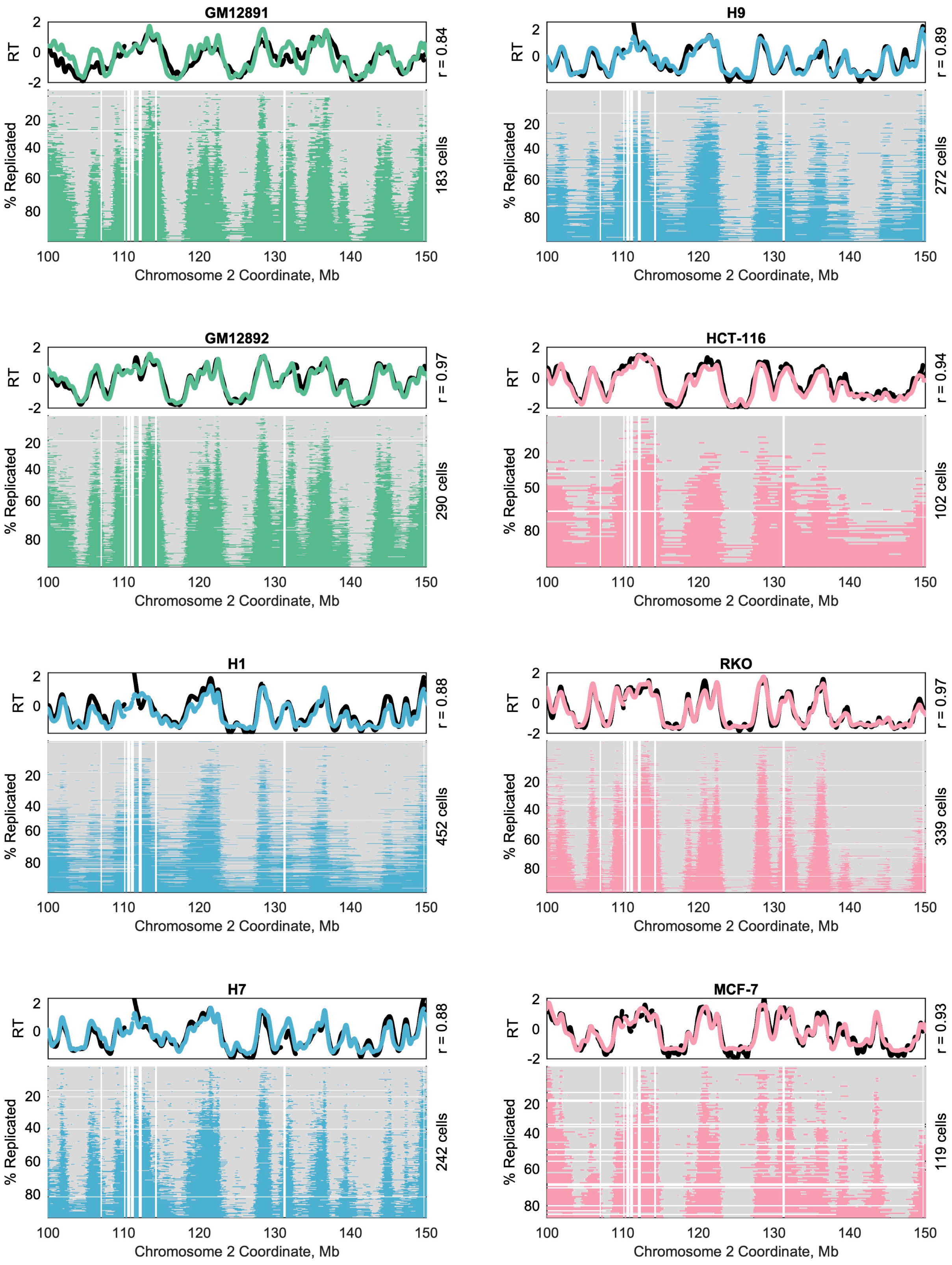
GM12891
r = 0.84
183 cells
H9
r = 0.89
272 cells
GM12892
r = 0.97
290 cells
HCT-116
r = 0.94
102 cells
H1
r = 0.88
452 cells
RKO
r = 0.97
339 cells
H7
r = 0.88
242 cells
MCF-7
r = 0.93
119 cells
RT
% Replicated
Chromosome 2 Coordinate, Mb

Supplementary Figure 7. **Replication state inference for thousands of single cells across human lymphoblastoid cell (green), embryonic stem cell (blue), and cancer-derived cell (pink) lines.** As in Figure 1e for all additional cell lines. Top panels: aggregate $S/G_1$ replication timing profiles from *in silico* sorting of single cells recapitulates the bulk-sequencing replication timing profile for the same cell line, in all cell lines. Bottom panels: single cell replication state in fixed 20kb windows. Each row represents a single cell.

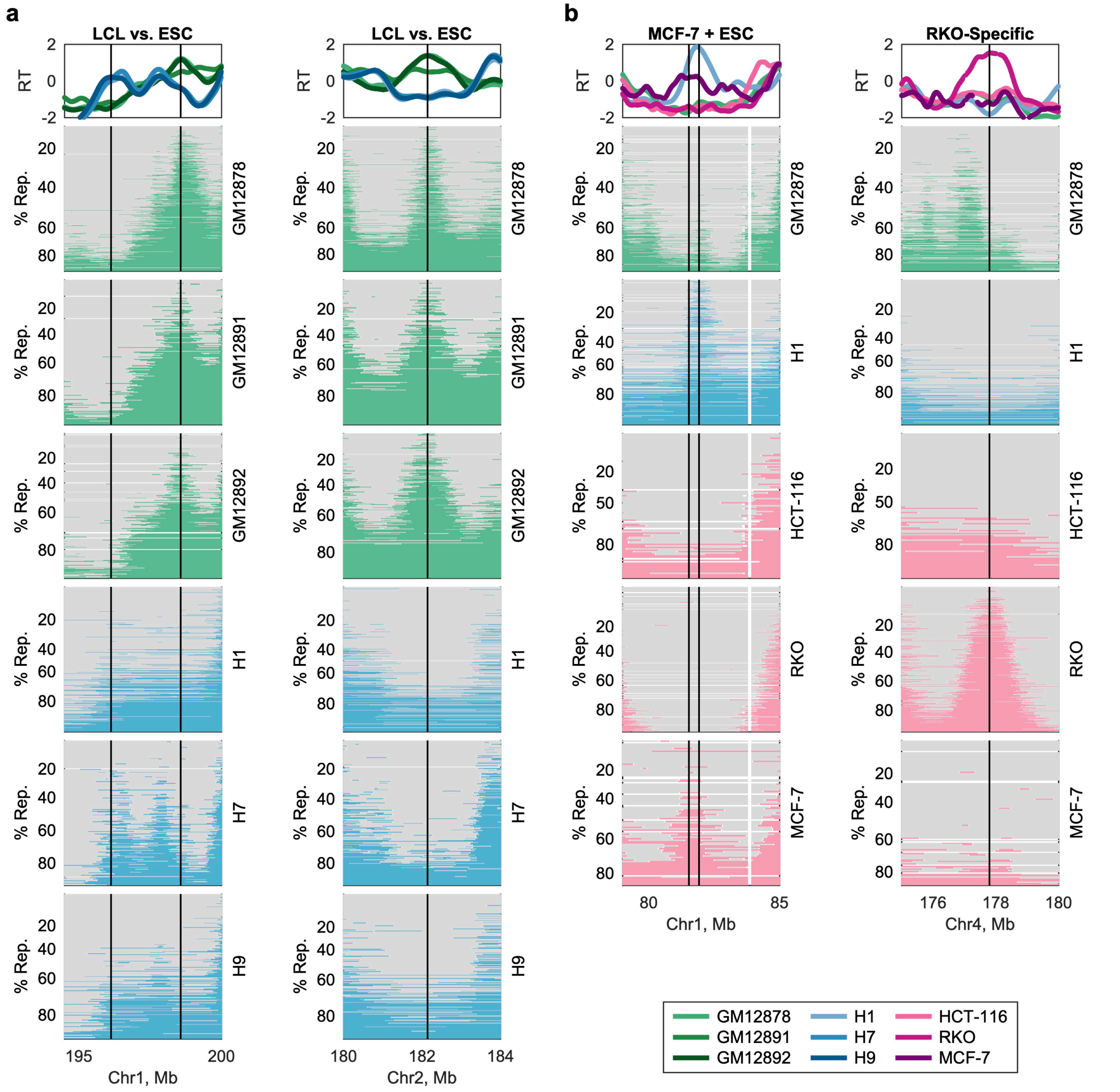


Supplementary Figure 8. **Cell-type- and cell-line-specific differences in the aggregate replication timing profiles are reflected at the single cell level.** The replication timing profiles shown are S/$G_1$ bulk-sequencing profiles. **a** Cell-type differences in replication timing between LCL and ESCs are observed at the single cell level. Left column: an ESC-specific peak in the bulk profile ~196.1Mb corresponds to a region of consistent replication initiation across all three ESCs, which is absent in GM1278 and GM12892 and fires sporadically in GM12891. Likewise, an LCL-specific peak in the bulk profile ~198.5Mb corresponded to a region of consistent replication initiation in LCL but not in ESC. Right: a second example of an LCL-specific region of replication initiation. **b** Differences in replication timing unique to individual cancer cell lines are consistent at the single-cell level. Left column: an MCF-7-specific peak ~81.5Mb is observed at the single cell level only in that cell line. To clarify that this initiation site is *not* observed in H1, a neighboring peak in ESC ~81.9Mb is also indicated with a vertical black line. Right column: an RKO-specific peak in the bulk profile is observed to be unique to RKO single cells as well.

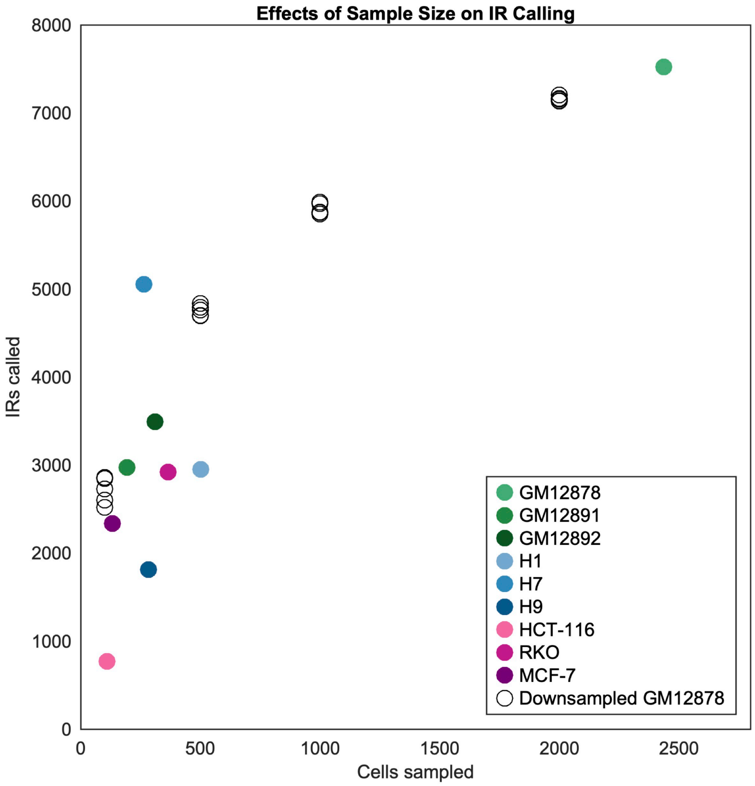


Supplementary Figure 9. **The number of initiation region (IR) calls increases non-linearly with increasing number of cells analyzed.** GM12878 cells were down-sampled (n = 250; 500; 1,000; 2,000) and initiation regions were called from each sample (black circles). Down-sampling was performed five times for each sample size to account for stochastic effects of small sample sizes. The observed number of IRs in each cell line is shown for comparison (colored dots).

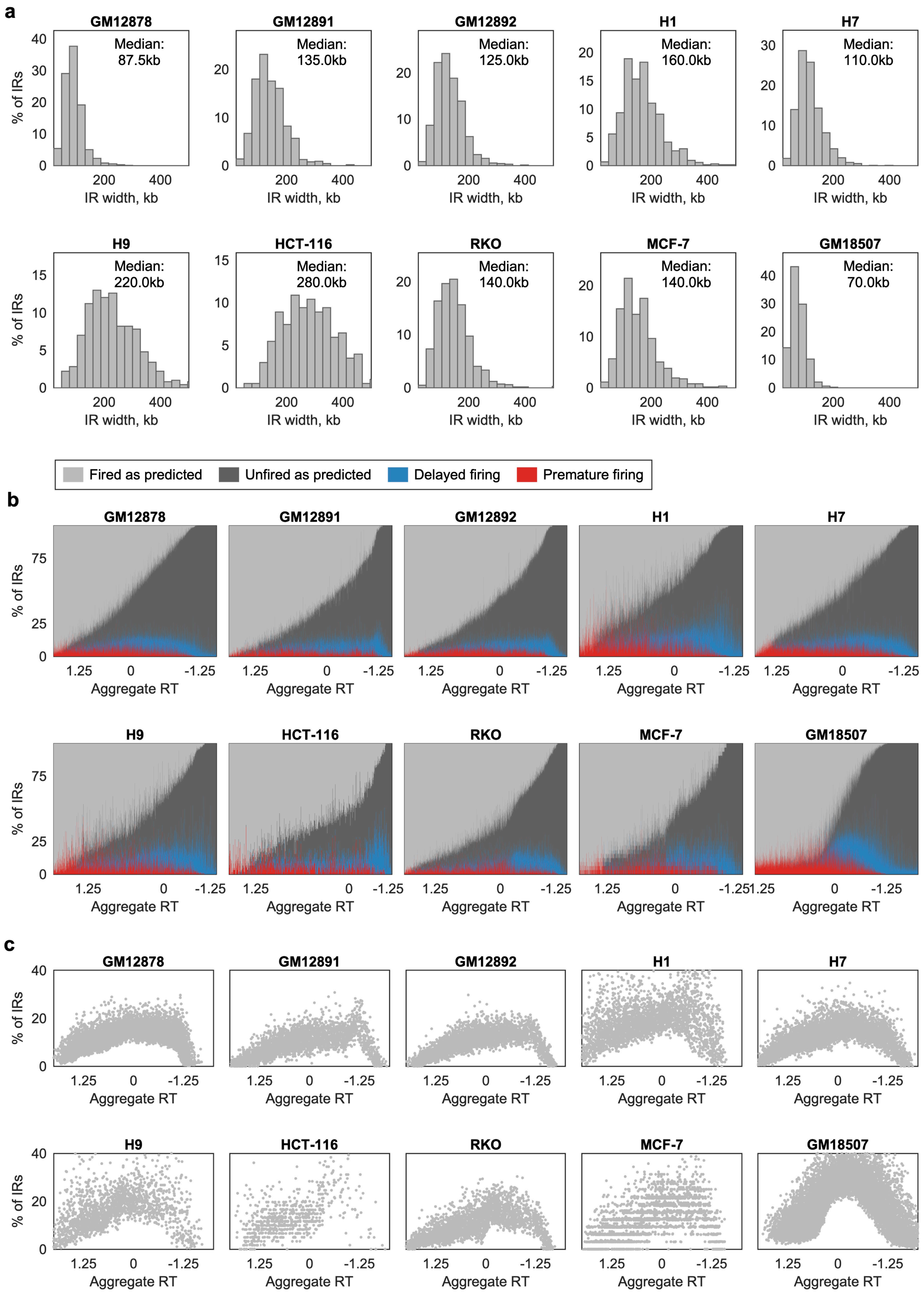
a
GM12878
Median: 87.5kb
GM12891
Median: 135.0kb
GM12892
Median: 125.0kb
H1
Median: 160.0kb
H7
Median: 110.0kb
H9
Median: 220.0kb
HCT-116
Median: 280.0kb
RKO
Median: 140.0kb
MCF-7
Median: 140.0kb
GM18507
Median: 70.0kb
% of IRs
IR width, kb
Fired as predicted
Unfired as predicted
Delayed firing
Premature firing
b
GM12878
GM12891
GM12892
H1
H7
H9
HCT-116
RKO
MCF-7
GM18507
% of IRs
Aggregate RT
c
GM12878
GM12891
GM12892
H1
H7
H9
HCT-116
RKO
MCF-7
GM18507
% of IRs
Aggregate RT

Supplementary Figure 10. **Initiation regions (IRs) fire in a similar, but not fixed order across cell lines. a** The distribution of IR widths is shown for all IRs genome-wide in each cell line under study. With the exception of HCT-116, where only 758 larger IRs were called, the distribution of IR widths was similar across cell lines and cell types, despite the noted cell-type specific difference in IR location and the ~10x reduction in sample size relative to GM12878. **b** IRs primarily fire within the expected order (light and dark gray), but sometimes fire earlier (red) or later (blue) than expected by a strictly determined ordering. There are no observed cell-type-specific differences. Compare to Figure 4c. **c** Variability in IR firing order is not uniform across S phase, but least pronounced at the beginning and end of S phase. This trend is not apparent in HCT-116 where almost all called IRs were early firing. Compare to Figure 4d. GM18507 cells were sequenced after DLP+ library preparation[2].

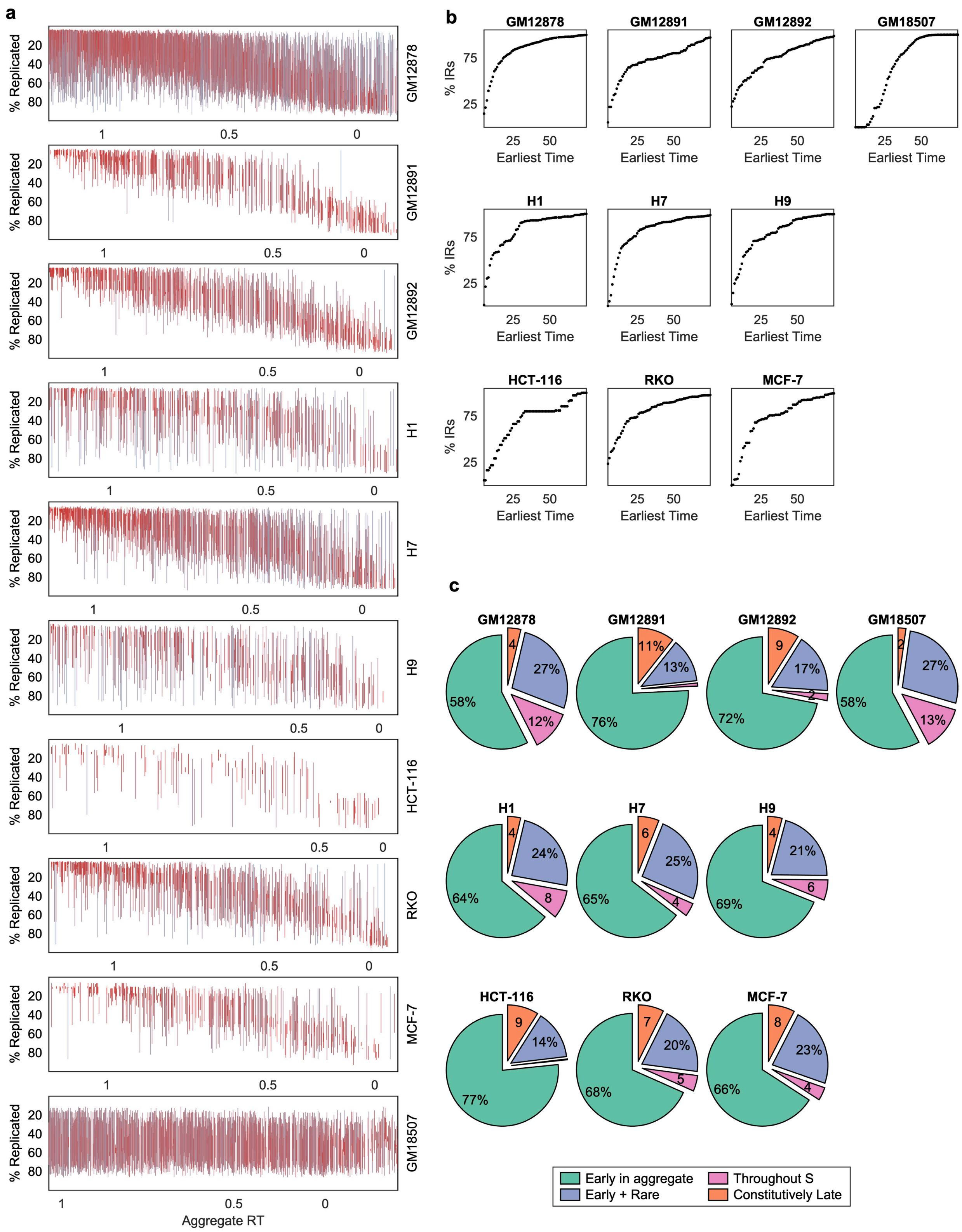

a
% Replicated
20
40
60
80
1
0.5
0
GM12878
GM12891
GM12892
H1
H7
H9
HCT-116
RKO
MCF-7
GM18507
Aggregate RT
b
GM12878
GM12891
GM12892
GM18507
H1
H7
H9
HCT-116
RKO
MCF-7
% IRs
75
25
25
50
Earliest Time
c
GM12878
4
27%
12%
58%
GM12891
11%
13%
76%
GM12892
9
17%
2
72%
GM18507
2
27%
13%
58%
H1
4
24%
8
64%
H7
6
25%
4
65%
H9
4
21%
6
69%
HCT-116
9
14%
77%
RKO
7
20%
5
68%
MCF-7
8
23%
4
66%
Early in aggregate
Early + Rare
Throughout S
Constitutively Late

Supplementary Figure 11. **Initiation regions (IRs) with early replication timing in aggregate tend to complete replication in early S phase, whereas IRs with late replication timing in aggregate tend to fire across a wider range of S phase. a** The range of IR firing was more constrained in IRs with early aggregate replication timing and less constrained in IRs with late aggregate replication timing. With a smaller sample size, we were able to call fewer IRs and we were able to corroborate the early S-phase firing of many IRs in a second cell. This produces a more constrained range in the additional cell lines (most pronounced in HCT-116). A random subsample of ¼ of the IRs is shown for GM18507, where the large number of IRs called obscures the trend. **b** Most IRs were observed to fire in early S phase in at least one cell. Each panel shows the cumulative percent of IRs that fire earliest before a given time in S phase. Across cell lines, 80-97% of IRs were observed to first fire in a cell <50% replicated. **c** Distribution of late IR classes across the additional cell lines. Although the smaller sample sizes resulted in calling about half as many IRs as in GM12878, a class of IRs replicating throughout S phase was observed in almost every cell line. The effects of sampling are most pronounced in the inflated proportion of IRs categorized as "early". GM18507 cells were sequenced after DLP+ library preparation[2].

Supplementary Table 1. **DNA sequencing details for each sequencing library generated in this publication.** Replicates are derived from culture of independent cryogenic vials. Each library was sequenced one to three times, as indicated.

| Cell Line | Fraction | Rep. | Instrument | Cycles | Center |
|---|---|---|---|---|---|
| GM12878 | $G_1$ | 1 | NovaSeq 6000 | 2 × 100 | 10x Genomics |
| GM12878 | Early S | 1 | NovaSeq 6000 | 2 × 100 | 10x Genomics |
| GM12878 | Full S | 1 | NovaSeq 6000 | 2 × 100 | 10x Genomics |
| GM12878 | Late S | 1 | NovaSeq 6000 | 2 × 100 | 10x Genomics |
| GM12878 | $G_2$ | 1 | NovaSeq 6000 | 2 × 100 | 10x Genomics |
| GM12878 | Unsorted | 1 | NovaSeq 6000 | 2 × 100 | 10x Genomics |
| GM12878 | Unsorted | 2 | HiSeq X Ten | 2 × 150 | GENEWIZ |
| GM12878 | Unsorted | 3 | HiSeq X Ten | 2 × 150 | GENEWIZ |
| GM12891 | Unsorted | 1 | HiSeq X Ten | 2 × 150 | GENEWIZ |
| | | | HiSeq X Ten | 2 × 150 | GENEWIZ |
| GM12892 | Unsorted | 1 | HiSeq X Ten | 2 × 150 | GENEWIZ |
| | | | HiSeq X Ten | 2 × 150 | GENEWIZ |
| H1 | Unsorted | 1 | HiSeq X Ten | 2 × 150 | GENEWIZ |
| H1 | Unsorted | 2 | NextSeq 500 | 2 × 36 | Cornell BRC |
| | | | HiSeq X Ten | 2 × 150 | GENEWIZ |
| | | | HiSeq X Ten | 2 × 150 | GENEWIZ |
| H7 | Unsorted | 1 | HiSeq X Ten | 2 × 150 | GENEWIZ |
| | | | HiSeq X Ten | 2 × 150 | GENEWIZ |
| H9 | Unsorted | 1 | HiSeq X Ten | 2 × 150 | GENEWIZ |
| HCT-116 | Unsorted | 1 | HiSeq X Ten | 2 × 150 | GENEWIZ |
| | | | HiSeq X Ten | 2 × 150 | GENEWIZ |
| RKO | Unsorted | 1 | HiSeq X Ten | 2 × 150 | GENEWIZ |
| | | | HiSeq X Ten | 2 × 150 | GENEWIZ |
| MCF-7 | Unsorted | 1 | NextSeq 500 | 2 × 75 | Cornell BRC |
| | | | HiSeq X Ten | 2 × 150 | GENEWIZ |
| | | | HiSeq X Ten | 2 × 150 | GENEWIZ |
